# Twisting DNA by Salt

**DOI:** 10.1101/2021.07.14.452306

**Authors:** Sergio Cruz-León, Willem Vanderlinden, Peter Müller, Tobias Forster, Georgina Staudt, Yi-Yun Lin, Jan Lipfert, Nadine Schwierz

## Abstract

The structure and properties of DNA depend on the environment, in particular the ion atmosphere. Here, we investigate how DNA twist -one of the central properties of DNA- changes with concentration and identity of the surrounding ions. To resolve how cations influence the helical twist, we combine single-molecule magnetic tweezer experiments and extensive all-atom molecular dynamics simulations. Two interconnected trends are observed for monovalent alkali and divalent alkaline earth cations. First, DNA twist increases monotonously with increasing concentration for all ions investigated. Second, for a given salt concentration, DNA twist strongly depends on cation identity. At 100 mM concentration, DNA twist increases as Ba^2+^ < Na^+^ < K^+^ < Rb^+^ < Li^+^ ≈ Cs^+^ < Sr^2+^ < Mg^2+^ < Ca^2+^. Our molecular dynamics simulations reveal that preferential binding of the cations to the DNA backbone or the nucleobases has opposing effects on DNA twist and provides the microscopic explanation of the observed ion specificity. However, the simulations also reveal shortcomings of existing force field parameters for Cs^+^ and Sr^2+^. The comprehensive view gained from our combined approach provides a foundation for understanding and predicting cation-induced structural changes both in nature and in DNA nanotechnology.

## Introduction

The structure and conformational flexibility of DNA play a central role in biological processes, including DNA read out, processing, condensation, and repair.^1–6^ A central property of DNA is its helicity.^7,8^ At physiological conditions, DNA has a twist corresponding to about 10.5 base-pairs per turn^9^ and it is known to be modulated by environmental conditions such as temperature^10–12^ and the ion atmosphere.^13–15^ In particular salt concentration strongly affects the structure and stability of DNA, as cations are essential for screening the negative charges on the sugar-phosphate backbone.^16–18^ In addition, preferential ion binding of the cations at the backbone or the grooves can influence the stability and structure.^19^ Consequently, the characteristic properties of DNA depend not only on the salt concentration and valence of the ions but also on the ion type.^13,14,20–24^

Characterizing and predicting the response of DNA to environmentally-induced changes is of central importance to understand biological function and to utilize DNA in biotechnological applications.^25–28^ Despite the importance, a comprehensive experimental characterization and a detailed understanding of how salt concentration and identity alter DNA twist are still lacking.

In order to fill this gap, we combine high-resolution magnetic tweezer (MT) experiments and all-atom molecular dynamics (MD) simulations. Single-molecule MT experiments are powerful tools to dissect global conformational changes of DNA.^29–31^ In MT, DNA molecules are tethered between a flow cell surface and magnetic beads (Figure 1a). Using external magnets, MT directly control the linking number of DNA at the single-molecule level^5,29–31^ and enable precise determination of DNA twist,^12,23,24^ in solution, with high resolution, and without requiring enzymatic reactions or staining (Figure 1a,b).

**Figure 1:**
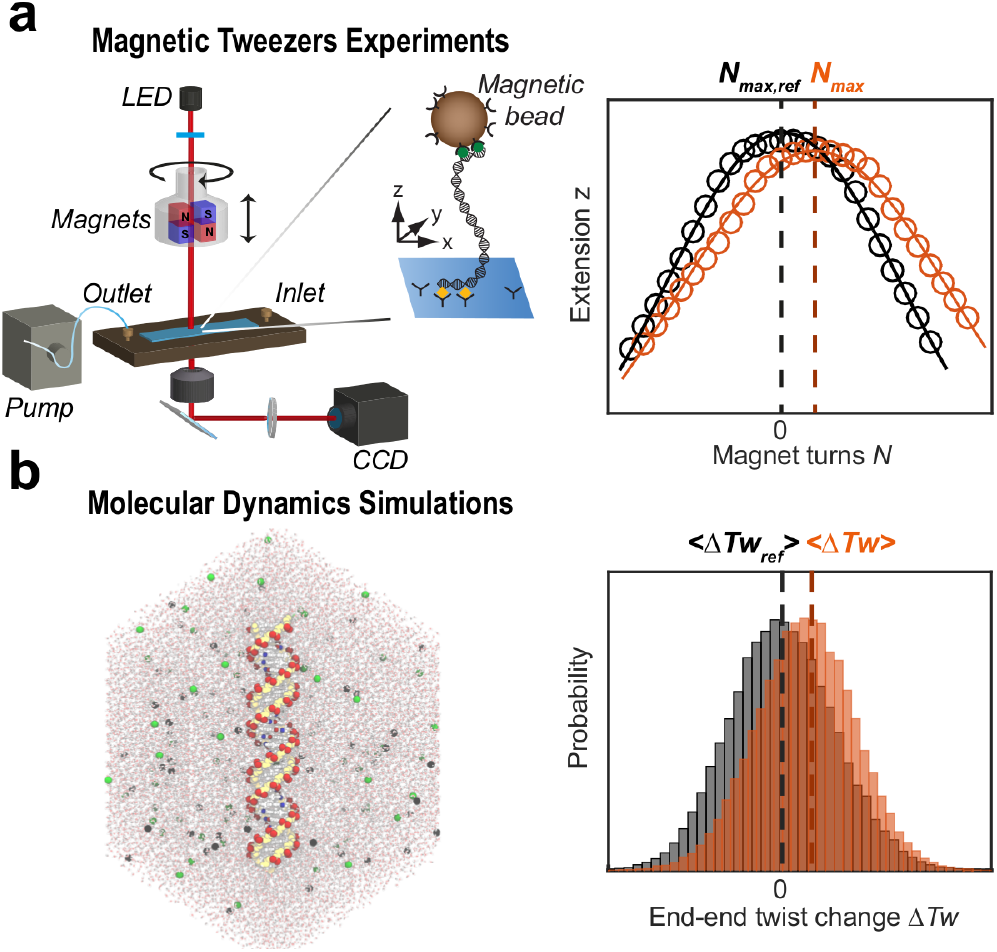
Magnetic tweezers and molecular dynamics simulations reveal the salt dependence of DNA twist. (a) Schematic of the magnetic tweezer set up with a flow cell that enables buffer exchange. The inset shows a DNA molecule tethered between the flow cell surface and a magnetic bead. Changes in DNA twist are experimentally determined from shifts in the maximum of extension vs. applied turns curves (symbols) that are fitted by Gaussian functions (solid lines). (b) Snapshot of an MD simulation showing the DNA, solvent, and the ionic atmosphere. Computationally, changes in twist are evaluated from shifts in the mean of the end-to-end plane twist. The data are for 100 mM KCl (the reference condition, black) and 1000 mM KCl (orange).

MD simulations are highly complementary to MT measurements, as they provide detailed, microscopic insights by resolving ion-nucleic acid interactions and the resulting conformational changes with atomistic resolution ^12,32–37^ (Figure 1c,d). The resolution afforded by MD simulations^38–43^ allows us to disentangle the contributions of backbone, nucleobases, ions, and water, which, in turn, provide a comprehensive view of the origin of ion specificity in DNA twist. Conversely, high-resolution twist measurements provide a useful test for current MD simulations. Changes in twist can be evaluated from simulations of moderately sized oligomeric DNA constructs and on the *µ*s-time scales accessible to state-of-the-art MD simulations. At the same time, DNA twist is a complex quantity that depends on the subtle interplay of multiple interactions. Therefore, DNA twist provides a stringent test for simulations, as was previously demonstrated for changes in twist with temperature^12^ and externally applied strain.^32,34^

Here, we report a comprehensive data set of DNA twist over a wide range of ion concentrations for the alkali and alkaline earth metal cations and quantitatively compare the experimental results to MD simulations. Our combined approach reveals two interconnected trends. First, DNA twist increases with increasing ion concentration. Our MD simulations show that increasing ion concentration leads to increased screening of the electrostatic repulsion between phosphate groups leading to a decrease in helical radius and crookedness,^44^ increase in sugar pucker, and ultimately increase in twist. Second, for a given salt concentration, DNA twist strongly depends on cation identity. The MD simulations explain the observed specificity through the preferential binding of the cations to the backbone and the nucleobase of the DNA. Ions with high backbone affinity (such as Li^+^) have a similar effect as large salt concentrations. They increase the sugar pucker, and twist and decrease the crookedness, and radius. In contrast, ions with intermediate backbone and nucleobase affinity (such as Na^+^) have a similar effect as low salt concentrations. They decrease the sugar pucker, and twist and increase the crookedness, and radius. While the MD simulations provide molecular level insights and correctly reproduce many of the experimentally observed effects, the results also reveal shortcomings and the need for further force field optimization for Cs^+^ and Sr^2+^.

## Methods

### Magnetic tweezers measurements

#### Magnetic tweezers instrument

We performed multiplexed single molecule magnetic tweezers experiments using a custom-build setup, described in detail previously.^45^ Precisely calibrated forces and torques are applied by a pair of permanent magnets (5 × 5 × 5 mm^3^, Supermagnete, Switzerland) arranged in vertical geometry^46^ with a gap of 1 mm and mounted in a holder that is controlled by a translation motor and a rotation motor. Motor control and real-time processing of the camera images was performed using a custom-written LabView program.^47^

#### Magnetic tweezers DNA construct and flow cell preparation

A 7.9-kbp DNA construct, prepared as described previously,^48^ was designed to allow torsionally constrained and oriented attachment to the bottom glass surface of a flow cell and paramagnetic beads. In brief, PCR-generated DNA fragments (≈600 bp) labeled with multiple biotin or digoxigenin groups were ligated to the DNA (7.9 kbp), to bind streptavidin-coated magnetic beads (1.0 *µ*m diameter MyOne beads, Life Technologies, USA) and the flow cell surface, respectively. To attach the DNA to the flow cell, the bottom coverslip was first modified with (3-Glycidoxypropyl)trimethoxysilane (abcr GmbH, Germany). Next, 75 *µ*l of a 5000x diluted stock solution of polystyrene beads (Polysciences, USA) in ethanol (Carl Roth, Germany) was dropcasted on the silanized slides, left to dry in a closed container, and baked at 80°C for 1 min. The polystyrene beads serve as fiducial markers for drift correction. A laser cutter was used to cut openings with a radius of 1 mm in the top coverslip for buffer exchange. The two coverslips were glued together by a single layer of melted Parafilm (Carl Roth, Germany), forming a ≈50 *µ*L channel that connects the inlet and outlet opening of the flow cell. After flow cell assembly, 100 *µ*g/ml anti-digoxigenin (Roche, Switzerland) in 1x PBS was introduced and incubated for 2 h. To reduce non-specific interactions of the DNA and beads with the flow cell surface, the flow cell was flushed with 800 *µ*l of 25 mg/ml bovine serum albumin (BSA; Carl Roth, Germany), incubated for 1 h and rinsed with 1 ml of 1x PBS. Next, the DNA construct was attached to streptavidin coated beads by incubating 0.5 *µ*l of picomolar DNA stock solution and 2 *µ*l My- One beads in 250 *µ*l 1x PBS (Sigma-Aldrich, USA) for 5 min and subsequently introduced in the flow cell for 5 min to allow formation of digoxigenin-anti-digoxigenin bonds. Subsequently, the flow cell was rinsed with 2 ml of 1x PBS to flush out unbound beads.

#### Magnetic tweezers twist measurements

Suitable double-stranded DNA tethers were identified as follows. The presence of multiple tethers was evaluated by rotating the external magnets to introduce negative supercoiling under high tension (*F* ≥ 5 pN), where for single DNA tethers the formation of plectonemes at negative linking differences is suppressed due to melting and the extension remains unchanged. In contrast, if multiple tethers are attached to the same bead, negative supercoiling results in braiding, which decreases tether extension. To assess whether DNA tethers were torsionally constrained, positive linking differences are introduced at low force (*F* = 0.4 pN), which results in plectoneme formation and a corresponding decreasing DNA extension for torsionally constrained tethers. In nicked DNA tethers, no linking difference can be introduced, and the extension remains constant on magnet rotation. Beads bound by multiple tethers or nicked tethers are discarded from further analysis. Following bead selection and testing, the buffer in the flow cell was exchanged for a buffer comprising 100 mM KCl and 10 mM Tris-HCl (pH = 7.0) using a peristaltic pump (flow rate ≈ 150 *µ*l min^*−*1^). Buffer exchange at this point is important from a practical view: we found that the direct substitution of PBS buffer for a buffer with certain salts, such as CaCl2, can induce irreversible sticking of the tethered beads. In addition, we use the buffer condition with 100 mM KCl, 10 mM Tris-HCl, pH 7.0, as the reference condition throughout. We recorded extension-rotation curves by changing the magnet rotation, and thus the linking number Δ*Lk*, in steps of two turns at low force (F = 0.5 pN). At each number of applied turns, the extension of the tether is recorded for 30 s (at a camera rate of 58 Hz), and the mean extension is calculated. Similar extension-rotation measurements were then carried out for a large range of ionic conditions. We investigated alkali chloride salts (LiCl, NaCl, KCl, RbCl, and CsCl; Sigma-Aldrich) at concentrations of 50 mM, 100 mM, 250 mM, 500 mM, and 1000 mM, and alkaline earth chloride salts (MgCl_2_, CaCl_2_, BaCl_2_, and SrCl_2_; Sigma-Aldrich) at concentrations of 1 mM, 2 mM, 5 mM, 10 mM, 25 mM, 50 mM, and 100 mM. All measurements used 10 mM Tris-HCl, pH = 7.0, as buffer. To ensure equilibration, we flushed a large volume of buffer at each new concentration through the flow cell (≥500 *µ*L or ≈10 cell volumes). During buffer exchange, we apply high force *F* = 6 pN, to ensure that the tethered beads do not rotate under the liquid flow and maintain constant linking number.

#### Magnetic tweezers data analysis

The linking difference Δ*Lk* at which the DNA is torsionally relaxed and the tether extension, therefore, maximal *N*max was determined by fitting the full extension-rotation curves with a Gaussian.^12^ The change in twist per base-pair at a given ion concentration and for each ion species is calculated from the difference of the relaxed state with respect to the relaxed state under reference conditions as ΔTw = 360°(*N*_max_ − *N*_max,ref_)*/*7900. For each condition, we measured at least two biological repeats and the reported values and error bars are the mean and standard deviations from at least 7 DNA molecules. Data processing and analysis were performed using custom routines in Matlab R2015a (MathWorks).

### Molecular dynamics simulations

#### Simulation setup

Our simulation system consists of a DNA duplex with 33 base pairs (bp), ions, and water. The sequence was 3’- GAGAT-GCTAA-CCCTG-ATCGC-TGATT-CCTTG-GAC-5’, identical to the one in Ref.^12^ To closely match the experiments, we simulated the monovalent ions Li^+^, Na^+^, K^+^, and Cs^+^ at bulk concentrations of 100, 250, 500, and 1000 mM and the divalent ions Mg^2+^, Ca^2+^, Sr^2+^, and Ba^2+^ at 25, 50, and 100 mM. The parmbsc1^49^ force field was used because exhaustive simulations have validated its ability to reproduce average experimental structures^50^ and have ranked it as one of the best in describing conformational dynamics of DNA.^33,51^ For the metal cations, we use the recently optimized force field parameters by Mamatkulov and Schwierz ^52^ and their extension^39^ for Ca^2+^. The choice of ion force field was motivated by the fact that the optimized parameters yield accurate ion-pairing properties as judged by comparison to experimental activity coefficients and accurate exchange kinetics as judged by experimental water exchange rates. In particular, the parameters were shown to resolve the fine differences between the distinct metal cations. ^38^ The force field parameters and charge density of the ions are listed in Table S3. Since Rb^+^ was not included in the ion parametrization,^52^ it was not simulated in this work. All systems were simulated for 3 *µ*s for monovalent and 5 *µ*s for divalent ions. We discarded the first 200 ns for equilibration and calculated the helical properties with `3DNA`^53^ and `do_x3dna`^54^ software packages and in-house scripts. Errors were estimated from block averaging with blocks of 200 ns. The cation distributions were analyzed using the canion tool of the software `Curves+`,^41^ and groma*ρ*s.^55^ Further details are in the Supporting Information.

#### Definition of helical twist

The definition of the helical twist is important for quantitative comparisons to experiments.^12,56–58^ We calculated the change of twist with three different methods: the sum of the local helical twist along the helix, the sum of the bp-twist contributions, and the end-to-end plane twist, where a local frame of reference is assigned to each end of the helix, and the change in twist is obtained by quaternion averaging (see Ref. ^12^ and Supporting Information for details). The results show that the absolute value of the calculated twist depends on the definition. However, the relative changes are similar for the three methods (Figure S1). In the following, we report the change in twist ΔTw obtained from the end-to-end plane measure. This definition is invariant to initial constant rotation offsets about the z-axis^12^ and closely mimics the experimental setup. As in experiments, we report the relative change in twist with respect to 100 mM KCl.

To evaluate the robustness of our results with respect to helix length, we repeated the calculations of ΔTw for progressively shorter subsequences by taking the central section of the helix while removing end bases from the analysis. The results show that using at least 15 bp provides consistent results (Figure S2). Therefore, the MD simulations can be compared to the MT results even though the helix length is much shorter in the simulations.

## Results and Discussion

We combine single-molecule magnetic tweezer (MT) experiments and extensive all-atom molecular dynamics (MD) simulations to resolve how salt concentration and identity of the ions influence the helical twist of DNA (Figure 1). In the following, we first report the experimental results, which reveal the changes induced by different metal cations. Subsequently, we provide a quantitative comparison to atomistic simulations and resolve the molecular mechanism of the conformational changes induced by different cations.

### Increasing concentrations of monovalent ions increase DNA twist

We systematically measured the DNA tether extension as a function of applied turns under a stretching force of *F* = 0.5 pN. In this low force regime, the response of tether extension is symmetric for over- and underwinding of the helix, and the DNA undergoes a buckling transition for both positive and negative applied turns.^29,31^ Past the buckling transition, the DNA forms positive or negative plectonemic supercoils, respectively, marked by a linear decrease of the tether extension with the number of applied turns.^29^ The rotation-extension curves reveal systematic shifts to positive turns with increasing ion concentration (Figure 2). We quantify the shifts by fitting the experimental data with Gaussian functions^12^ (Figure 1).

**Figure 2:**
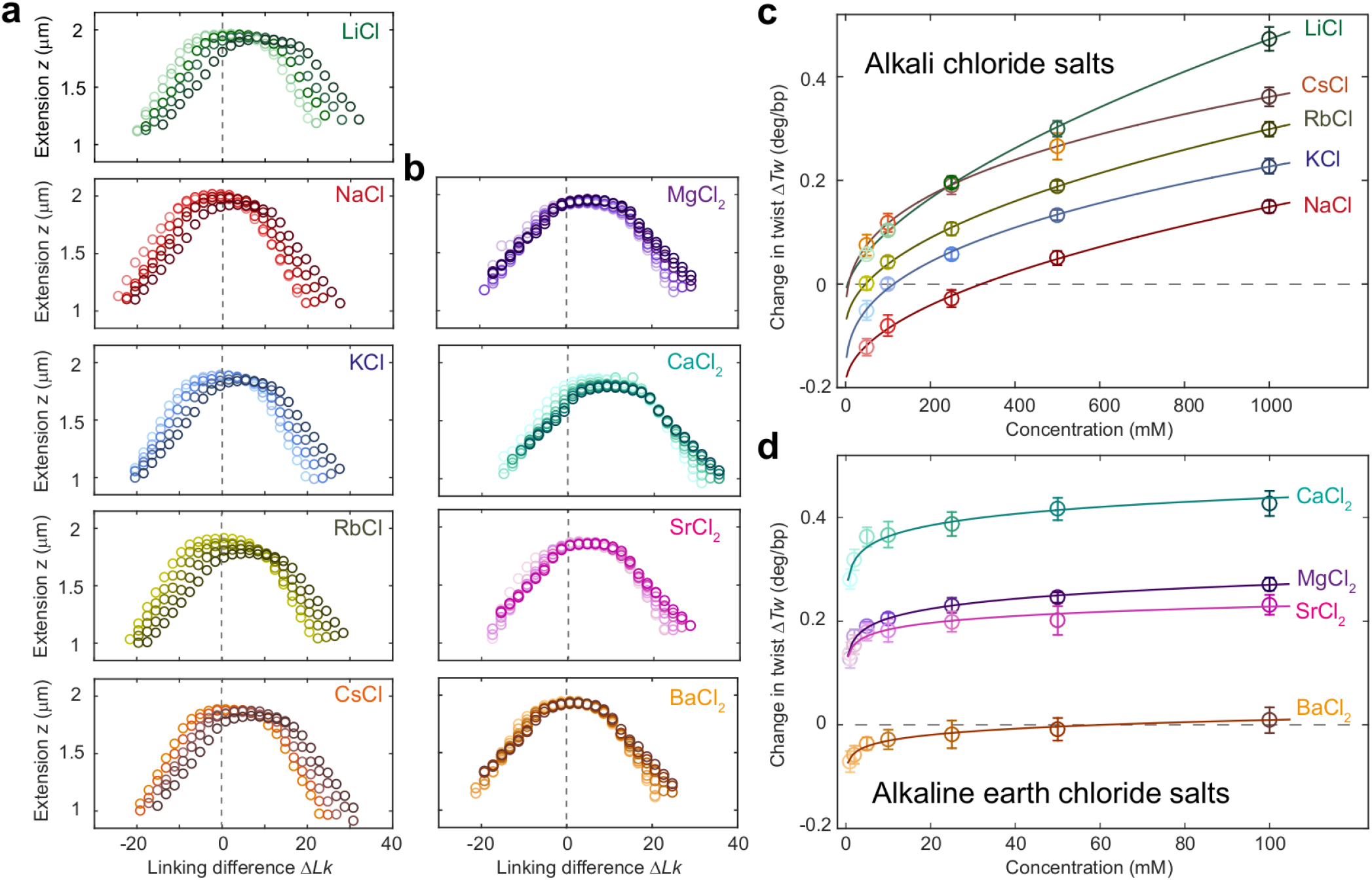
Magnetic tweezers measurements of the effect of salt on DNA twist. (a) DNA extension vs. rotation curves for alkali chloride ions at concentrations of 50, 100, 250, 500, and 1000 mM (light to dark data points). (b) DNA extension vs. rotation curves for alkaline earth chloride salts at 1, 2, 5, 10 25, 50, and 100 mM (light to dark data points). Zero Δ*Lk* (vertical dashed lines) corresponds to the point where the DNA is torsionally relaxed in 100 mM KCl. (c, d) Change in DNA twist with ion type and concentration for mono- and divalent metal cations. Symbols and error bars are the mean and standard deviations from at least seven independent measurements. Data are with respect to 100 mM KCl. Lines are fits to a power-law model (Equation 1; see main text).

Throughout, we used 100 mM KCl as our reference conditions. While the reference condition can in principle be selected arbitrarily, our choice of 100 mM KCl is motivated by the considerations that i) 100 mM monovalent ions correspond, approximately, to physiological ionic strength and intracellularly K^+^ is the most abundant cation, ii) 100 mM KCl is a condition that was consistently included in previous studies of the salt dependence of DNA twist using plasmids, ^13,14^ therefore, facilitating comparison, and iii) 100 mM KCl is readily and robustly accessible both in MT experiments and MD simulations.

A shift of the rotation-extension curves to positive turns corresponds to an increase in DNA twist with increasing ion concentration. The overall change in twist ΔTw going from 50 to 1000 mM salt concentration is approximately 0.3 °/bp or 1% for all monovalent ions (Figure 2a,c). The relative changes in twist with ion concentration are similar for all monovalent ions investigated and virtually identical for Na^+^, K^+^, Rb^+^, and Cs^+^ (Figures 2c and S3a). For Li^+^, DNA twist increases more rapidly with ionic concentration compared to the other monovalent ions.

Interestingly, the change in twist flattens with increasing salt concentration but does not saturate up to 1 M. In order to describe the change in twist with salt concentration, we tested two different empirical dependencies: i) a linear dependence on the log of the ion concentration as suggested previously^13^ in the concentration range 50-300 mM and ii) a power-law dependence. Fitting a linear dependence of the log of the ion concentration, we find slopes similar to the values reported by Anderson and Bauer,^13^ however, the fit does not capture the monovalent data very well (Figure S3b; reduced *χ*^2^= 29.1). In contrast, the data are much better described by a power-law dependence of the form

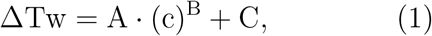

where c is the ion concentration and A, B and C are fitting parameters (Figure 2c, solid lines; reduced *χ*^2^ = 0.88). We find values for the scaling exponents B ≈ 0.5. Keeping B fixed and equal to 0.5, we find similar quality fits (Figure S3, reduced *χ*^2^ = 0.97). The complete set of fitted parameters are shown in Table S1.

In summary, DNA twist increases with increasing salt concentration. For monovalent ions, the behavior is well described by a scaling with the square root of the ion concentration. This scaling is in line with the dependence of the Debye length on ionic strength from simple Debye-Hückel theory.^59^

### DNA twist depends on monovalent ion- type

DNA twist depends significantly on the ion type, as evident from the vertical shift of the twist versus salt concentration curves for the different monovalent species (Figure 2c). In particular, Li^+^ is much more efficient at twisting DNA compared to the other monovalent cations. At 100 mM, the change of DNA twist is lowest for Na^+^ and highest for Cs^+^, with a difference of about 0.2 °/bp, similar to the change in twist induced by ion concentration. Therefore, ion concentration and ion type are equally important for determining DNA twist.

From the MT data, we can rank the cations according to their efficiency to induce DNA twist, i.e., to overwind DNA. A ranking of ions according to their effect on macroscopic properties was first introduces by Hofmeister,^60^ who ordered ions according to their efficiency to salt out proteins from solution. Ever since, a broad variety of rankings were found in systems ranging from simple electrolyte solutions to biological systems,^61–63^ including nucleic acids.^38,64,65^ In the generally accepted direct Hofmeister series, the order of the ions with respect to macroscopic properties roughly coincides with their charge density,^66^ while the opposite order is referred to as a reversed series.

At large salt concentrations, the change in twist follows a partially reversed Hofmeister series: Na^+^ < K^+^ < Rb^+^ < Cs^+^ < Li^+^, with the exception of Li^+^, the ordering corresponds to increasing twist with increasing ion size and thus decreasing charge density.

Comparing to previous data obtained by plasmid relaxation experiments combined with electrophoretic separation, both the ordering and trends with concentration determined by our magnetic tweezers experiments agree with the data by Anderson and Bauer,^13^ except for a slight deviation for Li^+^, which lies between K^+^ and Rb^+^ in their data. In contrast, Xu and Bremer observe essentially the same behavior for K^+^ and Cs^+^,^14^ in clear disagreement with both our and with the Anderson and Bauer data. However, it is worth noting that the emphasis of the Xu and Bremer study was on divalent metal ions, and they probed only a limited concentration range for monovalent ions.

Our results show that Li^+^ exhibits exceptional behavior as it gives rise to the largest twist and leads to a steeper increase of twist with concentration compared to the other monovalent ions (Figure 2c and S3). Anomalously large effects of Li^+^ were previously reported for ion counting experiment,^67–69^ nanopore translocation experiments,^70^ and SAXS studies,^71^ suggesting that Li^+^ ions interact more strongly with DNA, screen the negative backbone charge more effectively, and occupy a larger share of the ion atmosphere, compared to other alkali ions. The irregularity of Li^+^ has been suggested to result from its strong binding to the phosphate oxygens and expulsion from the nucleobases as previously reported for a dinucleotide system.^38^

In summary, DNA twist shows pronounced ion-specific effects, and the twist strongly depends on the type of metal cation. Larger monovalent ions overtwist DNA more strongly, except for Li^+^, which deviates from the overall trend and gives rise to the highest twist among the investigated monovalent cations. Taken together, the experimental data suggest that local interaction in addition to the overall composition of the ion atmosphere drives the changes of DNA twist.

### Divalent ions strongly modulate DNA twist

For divalent alkaline earth ions, DNA twist also increases monotonically with concentration and does not saturate up to 100 mM (Figures 2d and S4a). The monotonic increase obtained in our MT tweezers experiments (and MD simulations, see below) is in line with previous measurements^23^ for Mg^2+^, but contradicts the previously reported crossover from over- to underwinding observed in plasmid relaxation assays of Ca^2+^ and Ba^2+^ for increasing concentrations.^14^ The discrepancy might result from incomplete enzymatic relaxation of the plasmids at the highest divalent ion concentrations. Compared to the monovalent ions, divalent ions are much more efficient and the twist increase levels off already at much lower concentration. Fitting the dependence of DNA twist on divalent ion concentration, we find that in the range probed (1-100 mM), the data are equally well described by either a linear dependence of ΔTw on the log of the ion concentration or a power-law. The log of the ion concentration yields a reduced *χ*^2^ of 0.84 (Figure S4b); the power-law according to eq. 1 yields a reduced *χ*^2^ of 1.2 (Figure 2d). Here, the fitted scaling exponent B for different metal cations is *≈* 0.06 (see Table S2) and, therefore, much lower than for the monovalent species.

Similar to the monovalent ions, pronounced ion-specific effects are observed (Figure 2d). In the range 1-100 mM, the change in twist is lowest for Ba^2+^ and highest for Ca^2+^. The change in twist follows a partially reversed Hofmeister series: Ba^2+^ < Sr^2+^ < Mg^2+^ < Ca^2+^. With the exception of Mg^2+^, this ordering corresponds to decreasing twist with increasing ion size and thus decreasing charge density, which is opposite to the trend seen for monovalent ions.

Our results show that Ca^2+^ overwinds DNA significantly more (by 0.4 ^0^/bp at 10 mM) than any other mono- or divalent ion investigated in this study. The large effect of Ca^2+^ on DNA twist is in line with measurements by Xu and Bremer ^14^ and with previous reports that have used Ca^2+^ to induce positive supercoiling in plasmids.^72^

Except for Ca^2+^, the effect of the divalent ions on DNA twist follows the same ordering as in ion counting experiments that determine the occupancy in the overall ion atmosphere.^67^ However, the large differences in DNA twist for different divalent species are likely driven by specific local interactions and not simply due to the overall occupancy in the ion atmosphere.

### Molecular origin of increasing twist with increasing salt concentration

We performed extensive all-atom MD simulations to complement the MT experiments and provide a microscopic description of the ion effects on DNA. In the following, we first provide a quantitative comparison to the experiments for Na^+^ and K^+^, over the full concentration range (Figure 3a). We find that DNA twist increases with increasing concentration in the MD simulations and the results for Na^+^ and K^+^ are in quantitative agreement with experiments (Figure 3a).

**Figure 3:**
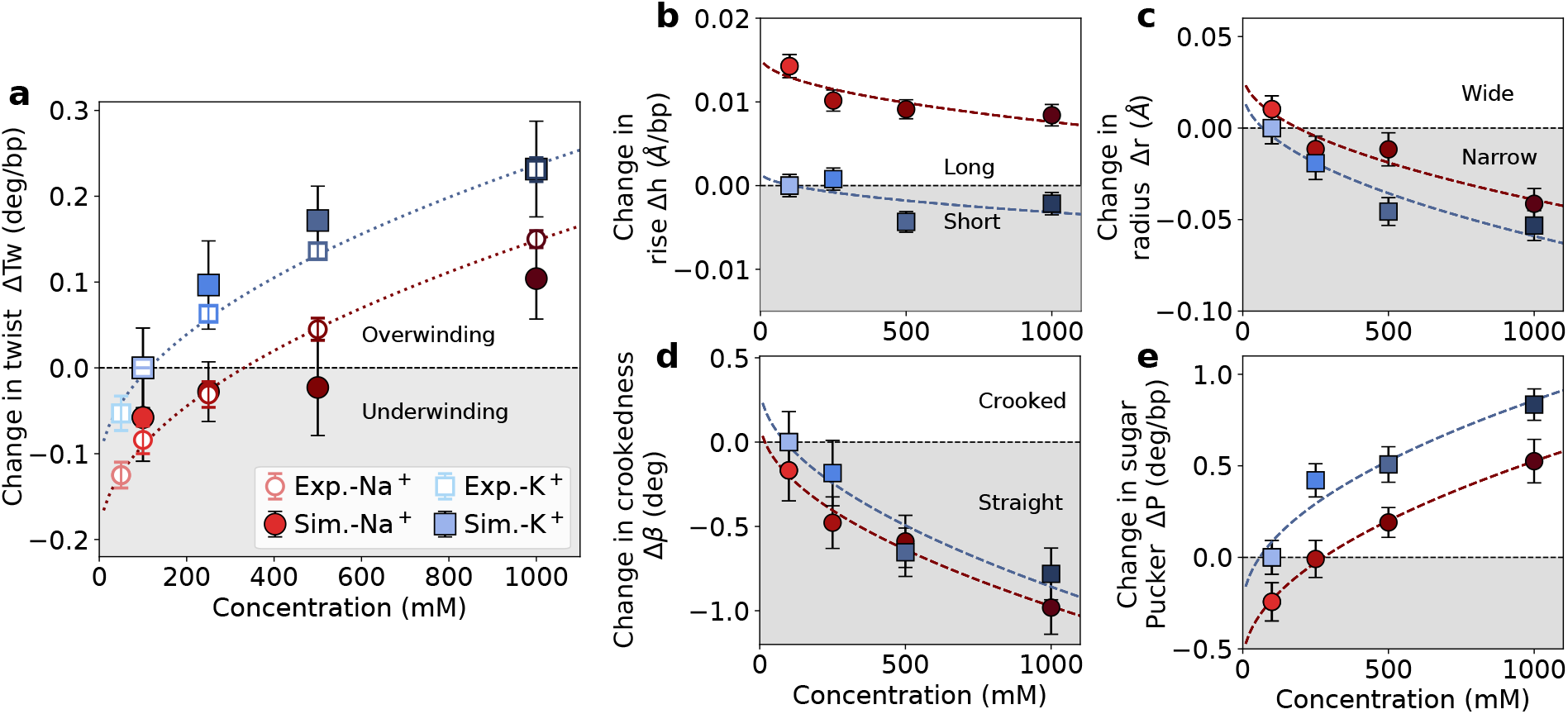
MD simulations reveal changes of DNA conformation with increasing Na^+^ and K^+^ concentration. (a) Quantitative comparison of DNA twist obtained from MT experiments (open symbols) and MD simulations (filled symbols) as a function of the ion concentration. Changes of the characteristic DNA properties as a function of the ion concentration: helix length *h* (b), radius *r* (c), crookedness *β*, (d), sugar pucker *P* (e). All parameters are reported relative to 100 mM KCl. Error bars correspond to standard errors obtained from block averaging. Dashed lines correspond to the fitting with the square root of concentration dependence. All fit parameters are listed in the Supporting Information.

Detailed molecular insight into the origin of this effect is obtained from characteristic DNA properties, including the helical rise, radius, crookedness, and sugar pucker as a function of ion concentration (Figure 3b-e). The helical crookedness^44^ *β* is determined from the ratio between the molecule’s extension *h* (i.e. the end-to-end distance) and the molecule’s length measured along the base pair centers *d* (i.e. the sum of distances between consecutive base pair centers) as cos *β* = *h/d*. Taken together, a consistent picture emerges: With increasing salt concentration, more and more cations adsorb at the backbone and compensate the negative charge (Figure 4h, i). In addition, an increasing amount of diffusive ions leads to increased screening. The resulting reduced electrostatic repulsion leads to a decrease in the DNA radius (Figure 3c). Helix narrowing is absorbed by a decrease of crookedness, an increase of the sugar pucker angle (Figure 3d, e) and a small decrease of the helical rise (Figure 3b). These changes result in increasing DNA twist with increasing salt concentration and are identical to the effects on DNA conformation that have been previously observed for the increase in DNA twist with decreasing temperature.^12,35^

**Figure 4:**
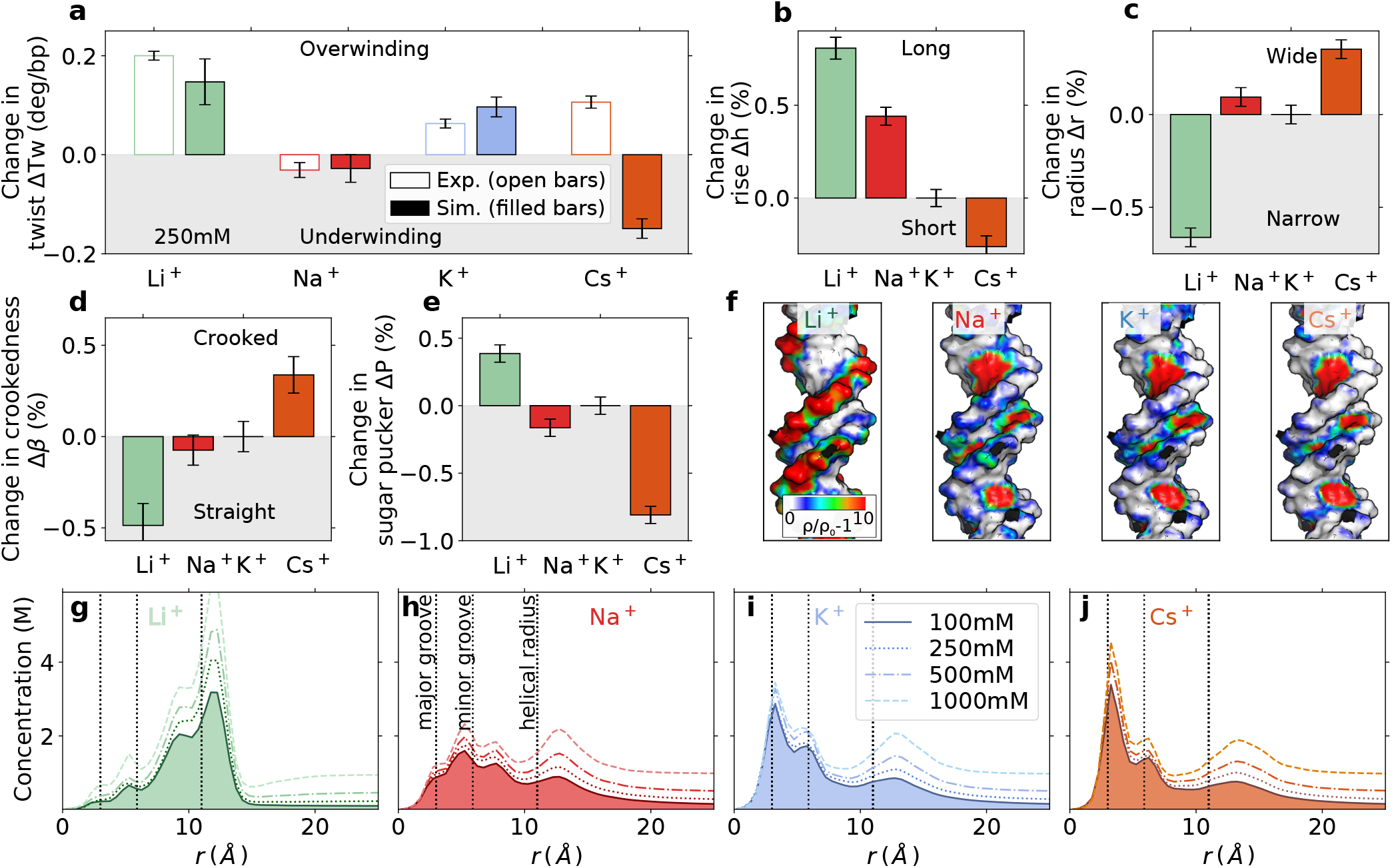
Origin of ion-specific effects on DNA twist for monovalent ions. (a) Quantitative comparison of change in DNA twist obtained from MT experiments (open bars) and MD simulations (filled bars) for 250 mM LiCl, NaCl, KCl, and CsCl with respect to 100 mM of KCl. Changes of the characteristic DNA properties for different monovalent cations: DNA helical length *h* (b), radius *r* (c), crookedness *β* (d), sugar pucker *P* (e). All results are relative to 100 mM KCl. Errors correspond to standard errors obtained from block averaging. (f) three-dimensional ion distributions obtained with groma*ρ*s^55^ and projected on the DNA surface. (g-j) Radial concentration profiles for 100 - 1000 mM bulk salt concentration.

### Origin of ion-specific DNA twist

Subsequently, we address the question of why DNA twist depends on the type of monovalent cation. A quantitative comparison of DNA twist obtained from MD simulations and experiments for different monovalent cations (Figures 4 and S1) finds excellent, quantitative agreement for Li^+^, Na^+^, and K^+^. However, the simulations predict underwinding for Cs^+^, in contrast to the experimental findings. This discrepancy reflects a shortcoming of the Cs^+^ force field due to an overestimation of the binding affinity of Cs^+^ to the nucleobases as will be discussed further below. Nevertheless, the results illustrate clearly that DNA twist strongly depends on the type of metal cation.

To elucidate the origin of the ion-specific effects, we calculated additional DNA properties (Figures 4b-e, S5 and S6). Overall, Li^+^, which induces the largest twist, simultaneously decreases the radius, and crookedness and increases the helical rise, and sugar pucker most strongly. By contrast, Cs^+^, which induces underwinding (compared to K^+^) in the simulations, increases the radius, and crookedness and decreases the helical rise, and sugar pucker. Overall, the influence of the ions on DNA twist, inverse radius, and sugar pucker in the simulations can be cast into a partially reversed Hofmeister series: Cs^+^ < Na^+^ < K^+^ < Li^+^, which deviates from the experimental data in the position of Cs^+^.

In general, ion-specific effects result from the distinct binding affinities of the cations to functional groups on biomolecules.^61,66,73^ Therefore, detailed insight into the origin of the Hofmeister ordering is obtained by resolving the preferred binding sites of the metal cations (Figure 4f-j). For simple nucleic acid systems, ions exhibit a high selectivity towards the individual binding sites.^38^ Ions with high charge density (such as Li^+^) preferentially bind to the phosphate oxygens on the backbone. In contrast, ions with low charge density (such as Cs^+^) preferentially bind to the N7 or O6 atoms of the nucleobases. ^38^

This trend is clearly reflected in the radial concentration profiles (Figure 4g-j) and the three-dimensional distributions (Figure 4f). The density of Li^+^ is highest at the backbone and significantly larger compared to Na^+^, K^+^, and Cs^+^. By contrast, the trend at the nucleobase (atoms N7 and O6) is exactly opposite. Here, the density of Cs^+^ is highest, followed by K^+^, Na^+^, and Li^+^ (Figure 4f-j), in agreement with previous work.^19,38,41^ Although individual ion-site affinities can explain most of the features of ion distributions, the spatial arrangement of the binding sites on the DNA influence further the preferential location of the ions. Cations can accumulate in specific volumes on the DNA, such as the minor and/or major groove (Figure 4f-j). For example, K^+^ has the highest density at the minor groove. In contrast, Li^+^ binds preferentially to the backbone, but it does not penetrate the minor groove. Overall, the ion distributions are determined by the interplay between site-specific binding affinities and steric effects.

Binding of the cations to the backbone or the nucleobases has opposite effects on DNA twist. Binding to the negatively charged phosphate oxygens on the backbone compensates the charges. The resulting diminished electrostatic repulsion decreases the DNA radius and increases the helical twist. Since Li^+^ has the highest backbone affinity, it gives rise to the highest twist among monovalent metal ions, in line with the experimental findings. On the other hand, binding to the nucleobases (major groove) has the opposite effect as neighboring bases along the helix are pulled together (Figure S5d), leading to a decrease in helical rise and to underwinding. Therefore, Cs^+^ reduces DNA twist in the simulations in contrast to the experiment. For intermediate cases, such as K^+^ and Na^+^, there is an interplay between nucleobase association and the screening of backbone charge repulsion.

The discrepancy between the twist obtained in simulations and experiments for Cs^+^ is likely caused by the overestimation of the affinity towards the nucleobases. More precisely, the binding affinity of Cs^+^ with the current force field is about 0.4 *k*_B_*T* higher at the O6 of the nucleobase compared to the phosphate oxygen. ^38^ Tiny differences in binding affinity can cause significant differences in the ionic distributions and, therefore, in the structure of DNA. In order to provide improvement, optimized Cs^+^ force fields are required that precisely reproduce the binding affinity of the cation toward the backbone and the nucleobases of DNA.

With the exception of Li^+^, the twist in the experiments increases with decreasing charge density for the monovalent ions (Figure 2c). Comparing K^+^ and Na^+^ in the simulations allows us to resolve the origin of the observed trend. Based on the ionic distribution profiles (Figure 4h,i), the density of K^+^ at the major groove (nucleobase) and the minor groove (around the phosphate groups of the backbone) is higher compared to Na^+^. Therefore, the two effects described above compete. The preferential binding of K^+^ at the major groove results in a reduced helical rise compared to Na^+^ (Figure 4b). Simultaneously, the strong location of K^+^ inside the minor groove leads to a more efficient screening of the negative backbone charges compared to Na^+^ resulting in, a smaller radius, larger sugar pucker angle, and larger twist (Figure 3a,c,e). This finding is further supported by the shrinking of the minor groove width (Figure S5b). The overall observed ion specific trends are therefore caused by a rather complex interplay of distinct binding affinities to the backbone and the nucleobases, as well as the accumulation of the ions in the major or minor groove due to steric effects.

### Mechanistic model for twisting DNA by metal cations

Our results show that metal cations influence the helical properties of DNA and small changes in the ionic environment lead to subtle conformational adjustments. The response of DNA to changes in the salt concentration and ion type is summarized in a simple mechanistic model (Figure 5). Our simulations show that the average sugar pucker angle and the twist are strongly correlated. Most importantly, the same correlation is obtained for different ion concentrations and ion types, suggesting a common origin of the overall changes (Figure 5a). Note that other properties, such as the sub-states of the backbone dihedral angles, which have been proposed to be correlated with local low- or high-twist states,^74,75^ did not show any significant changes across different concentrations for our systems, in agreement with recent observations^12^ (Figure S7).

**Figure 5:**
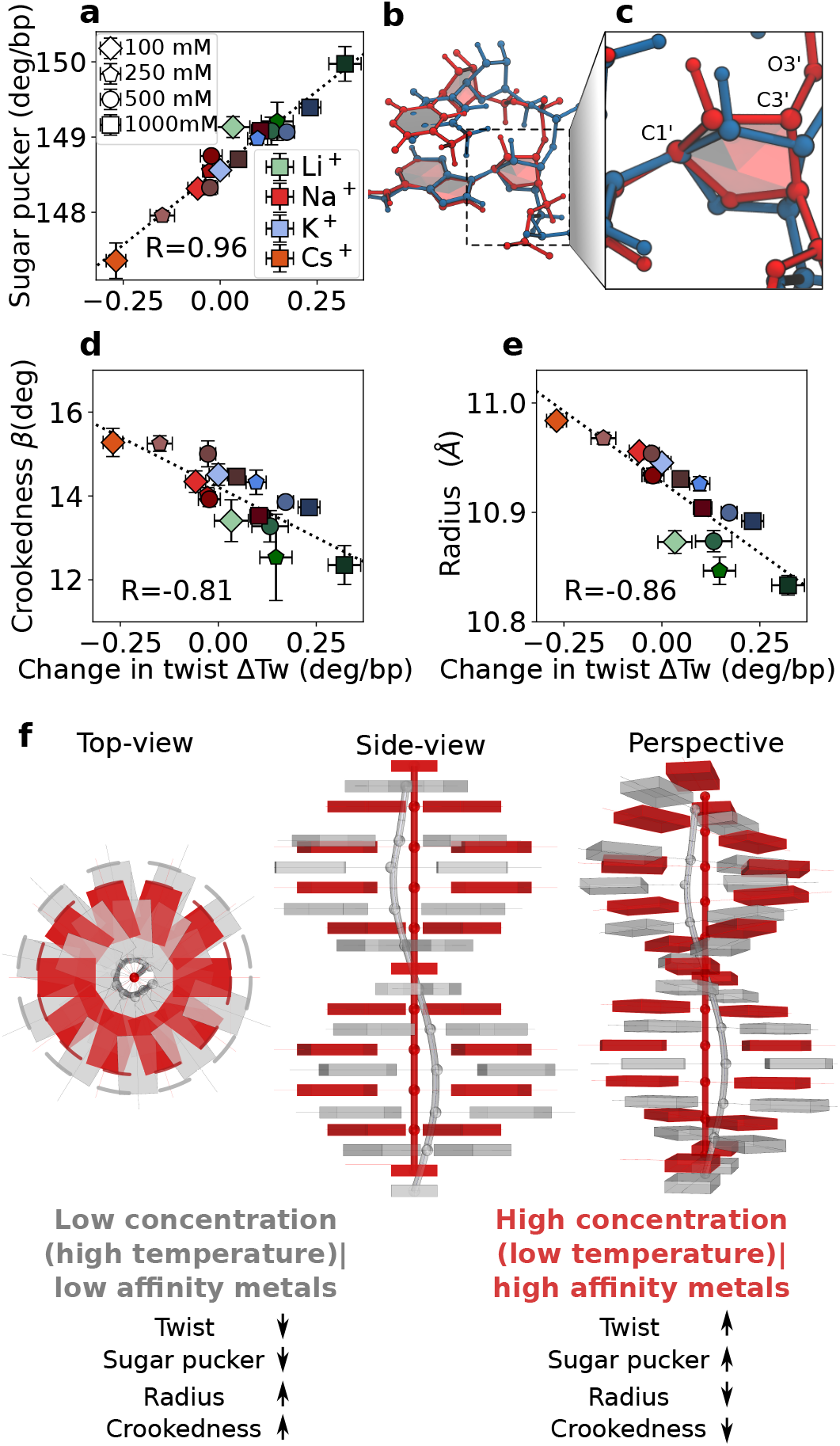
Mechanistic model for the changes of DNA structure with ion concentration and ion type. (a,d,e) Correlation between change in twist and sugar pucker angle (a), change in twist and crookedness (d), and change in twist and radius (e), for different ion concentrations and ion types. The reported value of R corresponds to the Pearson correlation coefficient. (b,c) Representative snapshots of two conformations with low (blue) and high sugar pucker (red). A moderate increase in the sugar pucker angle (from P=147° to P=157°) modifies the position of the phosphate group leading to a decrease in local radius (r=12.3 Å to r=9.7 Å), and crookedness (*β*=14.53° to *β*=6.28°), and an increase in local twist (Tw=33.3° to Tw=35.1°). (f) Simplified mechanistic picture: High salt concentrations and ions with high backbone affinity increase sugar pucker, and twist and decrease the radius, and crookedness (red structure). Low salt concentrations and ion with intermediate backbone and nucleobase affinity decrease sugar pucker, and twist and increase the radius, and crookedness (gray structure).

With the strong correlation between twist and sugar pucker, the conformational adjustments of the DNA in response to the ionic environment can be summarized as follows: The deoxyribose, that connects the nucleobase and the backbone of DNA, is flexible and adjusts readily to the ionic environment (Figure 5b,c) In particular, the sugar pucker adjusts depending on the repulsion between neighboring negatively charged phosphate groups along the backbone. Since the helical properties of DNA are coupled, changes in the sugar pucker affect the base-pair slide, and consequently, the radius of the helix.^34^ This effect is visualized in Figure 5b,c, which shows two DNA conformations: A moderate increase in the sugar pucker angle (from P=147° to P=157°) modifies the position of the phosphorous atoms leading to a decrease in radius (r=12.3 Å to r=9.7 Å), and crookedness (*β*=14.53° to *β*=6.28°), and an increase in twist (Tw=33.3° to Tw=35.1°). In this simplified mechanistic picture, high salt concentrations, ions with high backbone affinity (such as Li^+^), and low temperatures have the same effect. They increase the sugar pucker and twist and decrease the radius and crookedness (red structure in Figure 5f). Low salt concentrations, ion with intermediate backbone and nucleobase affinity (such as Na^+^), and high temperatures decrease the sugar pucker and twist and increase radius and crookedness (gray structure in Figure 5f). This model is further supported by the correlations between change in twist an crookedness, and with radius (Figure 5d,e). In this view, low salt concentrations and high temperature have the same effects on the twist, sugar pucker, radius, and crookedness,^12,35^ and also are known to destabilize the duplex, i.e., favor melting.^76^ Conversely, the increasing salt concentration increase DNA stability (melting temperature).^77^

### Importance of accurate force fields in predicting the twist for divalent metal cations

The influence of divalent cations on DNA twist is more complex than for monovalent ions. For divalent ions, ion binding leads to a local overcompensation of the backbone charge. Hence backbone segments can have positive or negative effective charges, leading to a complex interplay of attraction and repulsion as reflected in non-monotonic changes of DNA properties with increasing ion concentration in the simulations (Figures S8 and S9).

Figure 6a shows a quantitative comparison of the DNA twist obtained from simulations and experiments for different divalent cations at 50 mM salt concentration. For Mg^2+^ and Ba^2+^, the results of the MD simulations are in good agreement with experimental data, while the results for Ca^2+^ and Sr^2+^ significantly deviate from experiments, highlighting the sensitivity of the results on the ionic force field.

**Figure 6:**
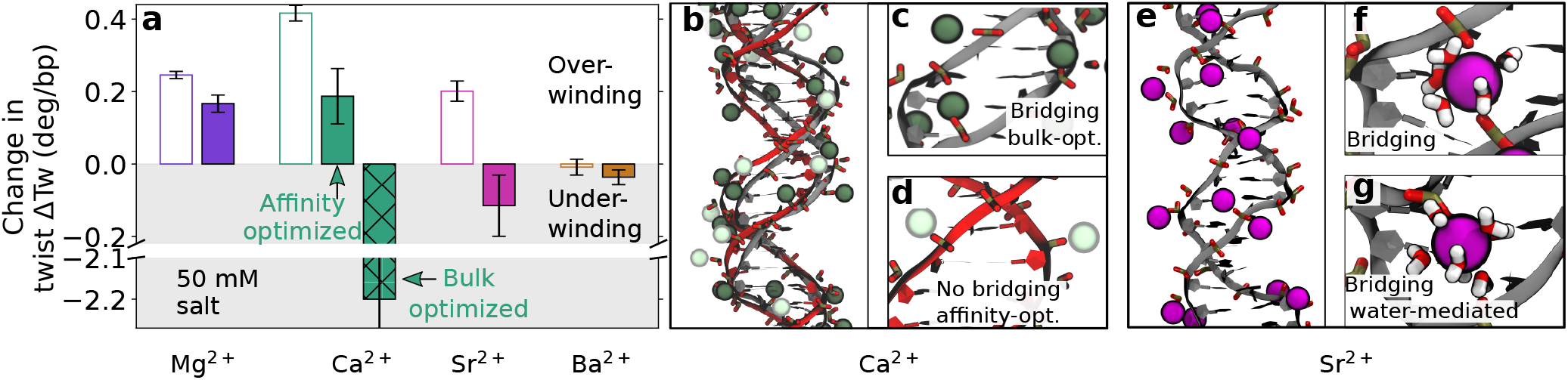
The effect of divalent metal cations on DNA twist. (a) Quantitative comparison of DNA twist obtained from MT experiments (open bars) and MD simulations (filled bars) for 50 mM MgCl_2_, CaCl_2_, SrCl_2_ and BaCl_2_. Data are with respect to 100 mM KCl and error bars correspond to standard errors obtained from block averaging. (b) Comparison of DNA structure obtained from simulations with bulk-optimized^52^ (gray) or (ii) affinity-optimized^39^ Ca^2+^ (red) force fields. (c) Spurious backbone bridging by Ca^2+^ ions with bulk-optimized force fields and undistorted structure with affinity-optimized force fields (d). (e) Simulation snapshot DNA structure with Sr^2+^ and formation of a backbone bridging (f) and a water-mediated backbone bridge (g).

Typically ion force fields are optimized to reproduce bulk properties and used in simulations without considering data related to interactions at specific nucleic acid sites.^78^ Here, we test the sensitivity of the twist using two available parameter sets for Ca^2+^: (i) bulk-optimized^52^ and (ii) affinity-optimized^39^ force fields. The first one was optimized based on ion bulk properties.^52^ The second one extends the applicability of the force field by including binding affinities of the cations toward the nucleic acids without changing ion bulk properties. ^39^

The bulk-optimized Ca^2+^ overestimates the binding affinity of the metal to the back-bone.^38,39^ As a consequence, in the simulations, Ca^2+^ ions bridge between phosphate groups on the backbone of the strands (Figure 6b, c), leading to distorted and under-twisted DNA structures. In contrast, the affinity-optimized force field^39^ significantly improves the agreement. Here, no backbone bridging is observed (Figure 6d) and the DNA is over-twisted in agreement with the experiments.

For Sr^2+^, bulk-optimized force fields predict direct and water-mediated backbone bridging and again lead to distorted and under-twisted DNA structures similar to Ca^2+^ (Figure 6e-g). Similar to Ca^2+^, improved force fields will be required for Sr^2+^ in order to enhance the agreement with experiments. Force field improvement will require further experimental data on the binding affinity of the metal cations in question to the phosphate oxygen and the nucleobases similar to the work by Sigel.^79^

## Conclusion

We have combined single-molecule magnetic tweezers experiments and all-atom molecular dynamics simulations to resolve how alkali and alkaline earth metal cations influence the helical twist of DNA. Two interconnected trends are observed: (i) DNA twist increases with increasing salt concentration, and (ii) DNA twist follows a partially reversed Hofmeister series for monovalent and divalent metal ions. At 100 mM, DNA twist increases as: Ba^2+^ < Na^+^ < K^+^ < Rb^+^ < Li^+^ *≈* Cs^+^ < Sr^2+^ < Mg^2+^ < Ca^2+^. Our results reveal that specifically adsorbed and diffusive ions reduce the electrostatic repulsion between the negatively charged backbone strands. The highly flexible sugar group readily adjusts to the resulting changes in the electrostatic environment, causing an increase of the sugar pucker angle and a decrease in radius and crookedness and, therefore, an increase of twist with increasing salt concentration (Figure 5a-c).

The ion-specific change in DNA twist and the resulting partially reversed Hofmeister series originate from the distinct binding affinities of the metals: Cations such as Li^+^, with high charge density, preferentially interact with the negatively charged phosphate groups on the DNA backbone, while ions such as Na^+^, with intermediate charge density, penetrate deeper into the grooves. Consequently, cations with high backbone affinity increase the sugar pucker, inverse radius, and twist, while metals with intermediate backbone and nucleobase affinity decrease the sugar pucker, inverse radius, and twist.

The overall changes in twist induced by different cations are subtle and provide a critical test of the accuracy of the ionic force fields. We find that for Ca^2+^ and, to a lesser extent, for Sr^2+^ bulk-optimized force fields^52^ lead to strong underwinding of the helix due to bridging between phosphate groups, in contrast to the experimental findings. Affinity-optimized force fields^39^ provide significant improvement. Still, the agreement with experiments for Ca^2+^ is not quantitative likely because the nucleobase affinity was not included in the force field optimization.

For Cs^+^, we similarly find that MD simulations underestimate the increase in twist induced by the cations, likely due to an overestimate of the Cs^+^ affinity to the nucleobase. Taken together, our results reveal the need for further force field optimization, in particular for Sr^2+^ and Cs^+^. Unfortunately, the calculation of DNA twist is computationally too expensive to test a broad parameter range to further optimize the cation force fields. Therefore, experimental data on the binding affinity for all metal cations to the phosphate oxygen and the nucleobases would be an invaluable contribution to the field and the next step in providing more accurate force fields for cations.^39,79,80^

While the changes in twist with ion identity and concentration are moderate, they would be amplified for long genomic DNA or large DNA assemblies. For example, the twist change induced by different types of divalent ions at constant ionic strength is similar in magnitude to the changes obtained by inserting or deleting base pairs in 3D origami assemblies with curvature.^27^ In summary, the results presented here serve as the baseline to critically test existing models of metal ion-nucleic acid interactions and provide a foundation to understand and predict changes in more complex nanostructures induced by the ionic environment.

## Supporting information

Experimental and computational details and supporting analysis

## Supporting Information

The Supporting Information is available free of charge on the publications website. It contains experimental and computational details and supporting analysis of the MT measurements and the MD simulation results.

## Acknowledgements

We thank Thomas Nicolaus for laboratory assistance and David Dulin for helpful discussions. N.S. and S.C.L. acknowledge financial support from the DFG (Emmy Noether program, Grant No. 315221747). LOEWE CSC and GOETHE HLR are acknowledged for supercomputing access. J. L. acknowledges financial support from the DFG via SFB863- Project-ID 111166240, A11.

