## Supplementary material for "Twisting DNA by Salt": Experimental and computational details and supporting analysis

### Supplementary methods

#### Molecular dynamics simulations

Independent simulations of the dsDNA for each ion type and salt concentration were performed using the Gromacs simulation package[S1] version 2018.1. Simulations included periodic boundary conditions and electrostatic interactions were treated using particle-mesh Ewald summation[S2] with tin foil boundary conditions. We used a 2 fs time step with constraints on the hydrogen bond atoms using the LINCS algorithm.[S3] The long-range electrostatic interactions were treated with cubic interpolation and a Fourier space grid of 0.12 nm. Lennard Jones interactions and close Coulomb real space interactions were cut-off at 1.2 nm. Errors from the truncation of LJ interactions were accounted for by long-range dispersion correction for energy and pressure.

**Simulation protocol**: The dsDNA structure was placed in an orthorhombic dodecahedral box, assuring a minimal distance of 1.5 nm to the edge. The simulation box was filled with 71011 TIP3P water molecules.[S4] Subsequently, water molecules were randomly replaced by ions to obtain a neutral system with the desired salt bulk concentration. A pre-equilibration protocol, consisting of energy minimization, NVT equilibration and NPT equilibration, was completed prior to production runs. During the pre-equilibration process, the heavy atoms of the nucleic acids were constraint with a soft harmonic potential with a force constant of 1000 kJ/(mol $nm^2$) to allow the equilibration of the solvent around the nucleic acid. Energy minimization used the steepest descent algorithm with a maximum of 50000 steps. Later, we employed 1 ns NVT and 1 ns NPT simulations to further equilibrate the system while keeping the positions restraints.

Finally, unrestrained runs were performed in the NPT ensemble. Trajectories were 3 $\mu$s long for monovalent cations and 5 $\mu$s long for divalent cations. NPT simulations used the isotropic Parrinello-Rahman barostat[S5] and the velocity rescaling thermostat with a stochastic term.[S6]

**Analysis of trajectories**: Helical structural parameters were analyzed with the broadly used software tools `3DNA`[S7] and `do_x3dna`,[S8] complemented with in-house scripts. Particularly, we obtained the helical-twist, helical-rise $h$, the base-pair (bp-)twist, and bp-rise $d$, the radius $r$, the sugar pucker $P$. r was defined as the mean distance of the phosphorous atoms to the helical axis. Finally, the helical crookedness $\beta$, previously introduced,[S9] is measured from the ratio between $h$ and the base-pair centers $d$ ($\cos\beta = \mathrm{h/d}$).

For the analysis of the helical parameters, the last three bases at each end were not considered. The reported error values correspond to the standard error of the mean (SEM) after block averaging. The cations distributions were analyzed using the `canion` tool of the software `Curves+`,[S10] which calculates the concentration profiles in untwisted helicoidal coordinates, allowing a detailed view of the interaction of the ions at specific volumes in the DNA. Complementary, groma$\rho$s was used to obtain the three-dimensional distribution of the ions,[S11] and the results were visualized with `pymol`.[S12]

#### Definitions of the twist in the simulations

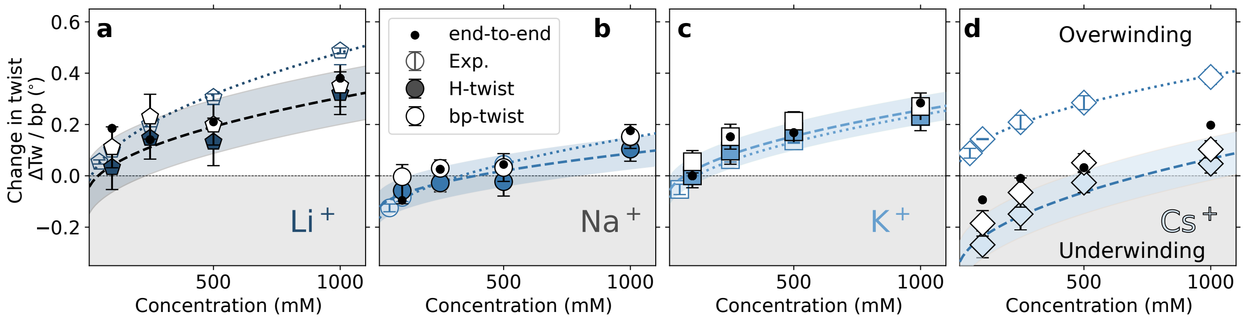


Figure S1: Comparison of ΔTw obtained from three different definitions: (i) the sum of the bp-twist (white symbols), (ii) the sum of the helical-twist (filled symbols), and (iii) the end-to-end plane definition twist as described by Kriegel et al.[S13] (points). Empty symbols are the experimental results. The dotted lines are the fitting of the experimental results to the square root of the cation concentration. To compare the different methods, the dashed line and the transparent area corresponding to the fitting of the helical-twist to the square root of concentration and its error are plotted.

To compare the structural changes of the DNA with the changes observed in experiments, we calculated the change in the helical twist using three different definitions, as previously discussed:[S13] (i) the sum of the bp-twist, that is, the sum of local twist of a dinucleotide step. (ii) the sum of the helical-twist. It corresponds to the sum of the rotation angles about the helical axis that brings successive base pairs into coincidence. (iii) the end-to-end plane definition twist described by Kriegel et al..[S13] In this definition, a base-fixed coordinate reference frame[S14] is assigned to both ends of the helix using quaternions averaging,[S15] and the total helical twist corresponds to the rotation of the X-Y plane of the reference frames.[S13] For each definition, we obtained the change in the twist for every recorded snapshot.

The three methods yielded similar relative changes for the change in twist $\Delta$Tw, as shown in Figure S1. In the following, we report the end-to-end twist because it has the property of being invariant with respect to initial constant rotation offsets about the z-axis,[S13] and it reproduces the closest the experimental setup where the initial torsional offset of the beads is not known. For a detailed discussion, see Ref.[S13]

#### Influence of duplex length on the calculated helical twist

Furthermore, recent experimental[S16] and computational evidence[S17,S18] have shown that the sequence strongly influences the equilibrium conformation of DNA. In our magnetic tweezers experiments, one would expect this effect to average out as measurements included a mixed sequence with 7900 bps. In contrast, for simulations of 33 bp it is not clear, a priori, how the sequence and length of the simulated helix may affect the relative changes in twist. To assess this effect, we computed $\Delta$Tw for different duplex lengths. Hereby, we subsequently removed the last bp at each end of the simulated helix (Figure S2). Our results show similar changes up to 15 bp (see Figure S2), which indicates that sequence effects average out with the 33 bp sequence used in the simulations.

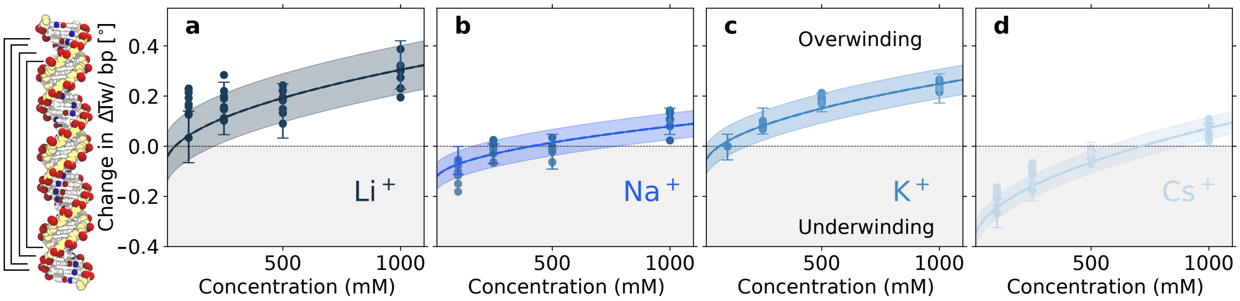


Figure S2: Dependence of ΔTw on the duplex length. ΔTw was calculated for the central section of the helix of different lengths between 15 and 27 bps by removing the bases at each end (scattered points). For comparison, the fit (continuous line) for the change in twist for 27 bps and its error (the transparent area) are shown.

### Supplementary experimental results

#### Fit of the experimental change in twist with concentration

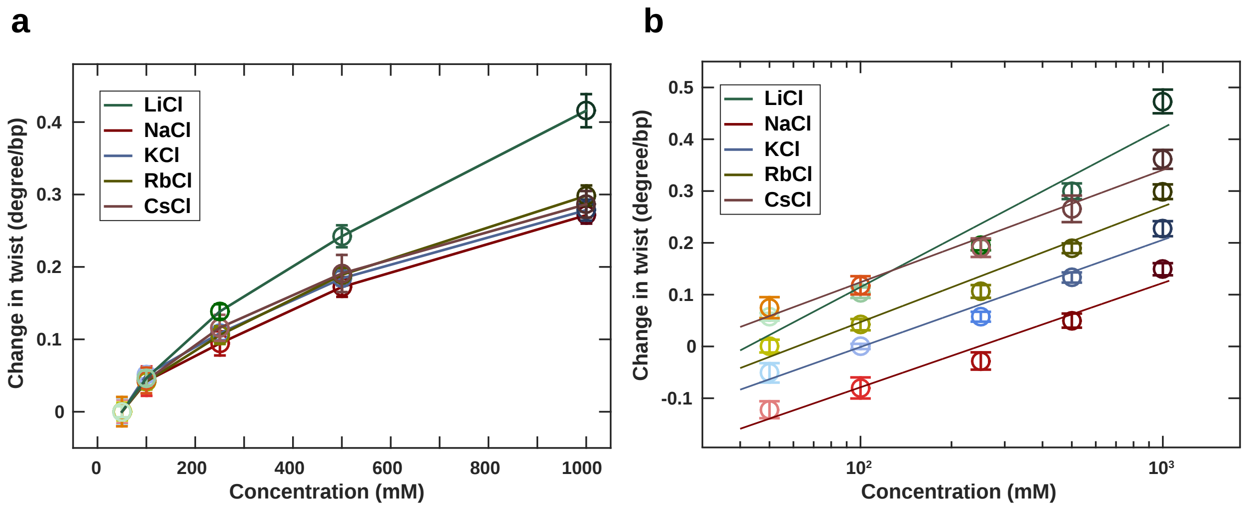


Figure S3: Change in DNA twist for monovalent cations as function of concentration. Symbols and error bars are the mean and standard deviations from at least eight independent molecules (except for $Cs^+$).(a) Data are shifted to the corresponding 50 mM data point and shown in a linear-linear plot. (b) Data on a log-linear plot with respect to 100 mM KCl. Lines in panel (b) are the fit to the log of the ion concentration model with a reduced $\chi^2 = 29.1$.

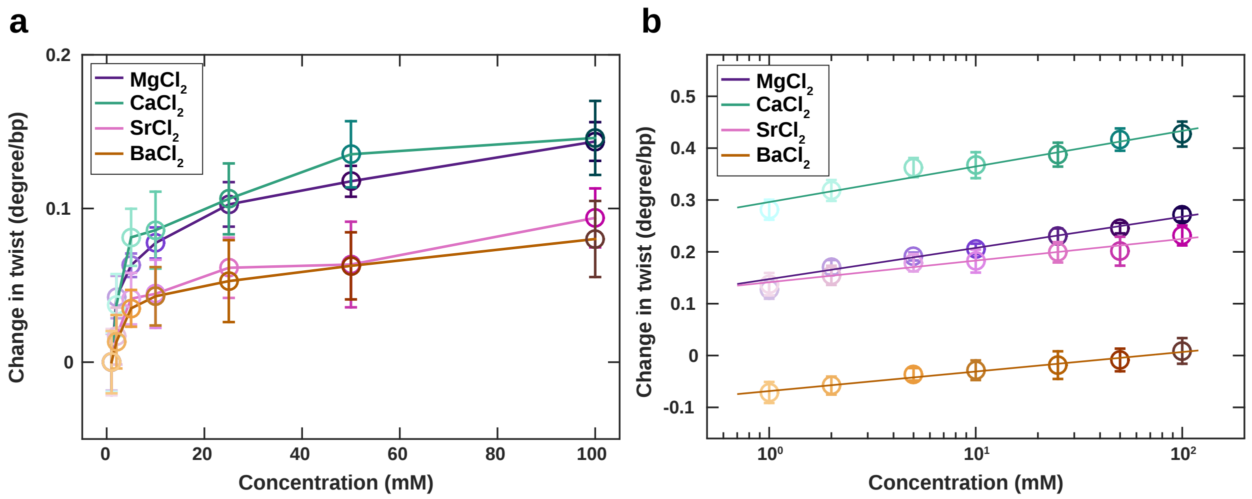


Figure S4: Change in DNA twist with ion type and concentration for divalent ions. Symbols and error bars are the mean and standard deviations from at least seven independent molecules. (a) Data shifted to the corresponding 1 mM data point and shown in a linear-linear plot. (b) Data plotted in a log-linear plot with respect to 100 mM KCl. Lines are the fit to the log of the ion concentration model with a reduced $\chi^2 = 0.84$.

### Supplementary simulation results

#### Changes in major and minor grooves

As shown in Figure S5, the width of the minor groove changes with the ion type and concentration. $Li^+$ compacts the minor groove the most, followed by $K^+$, $Na^+$, and $Cs^+$. In contrast, the major groove width is essentially unaffected by $Li^+$ and $Na^+$, but it is compressed by $K^+$ and $Cs^+$ (Figure S5), reflecting the accumulation of the ions in the grooves as discussed in the main text.

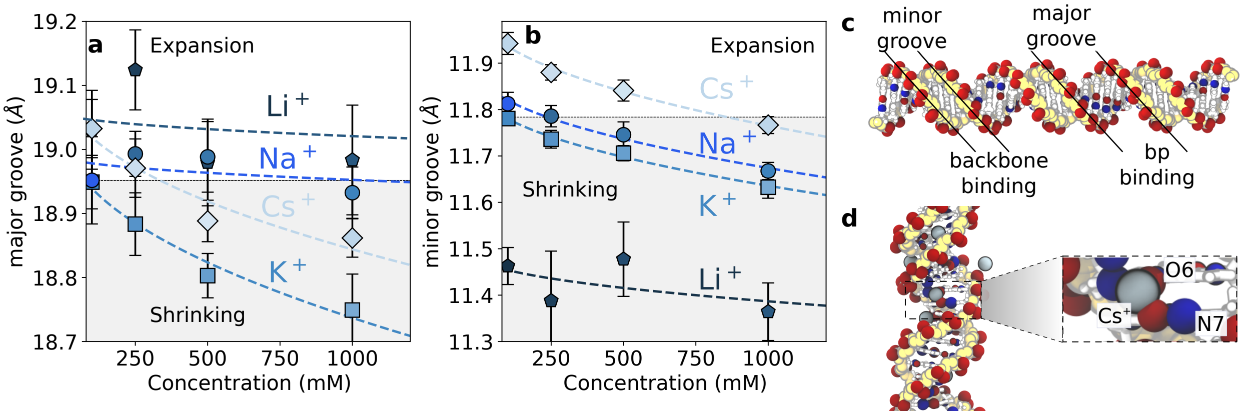


Figure S5: (a,b) Changes in the minor groove and the major groove width depending on ion type and concentration. Error bars correspond to the SEM. (c) Snapshot of the simulated DNA structure indicating the binding volumes. (d) Snapshot of s segment of DNA with a detail of $Cs^+$ bound at the major groove (atoms N7 and O6) pulling two consecutive bases together.

#### Helical parameters as function of ion type and concentration

Similar to ΔTw, we defined the change for all the helical properties relative to 100 mM of KCl.

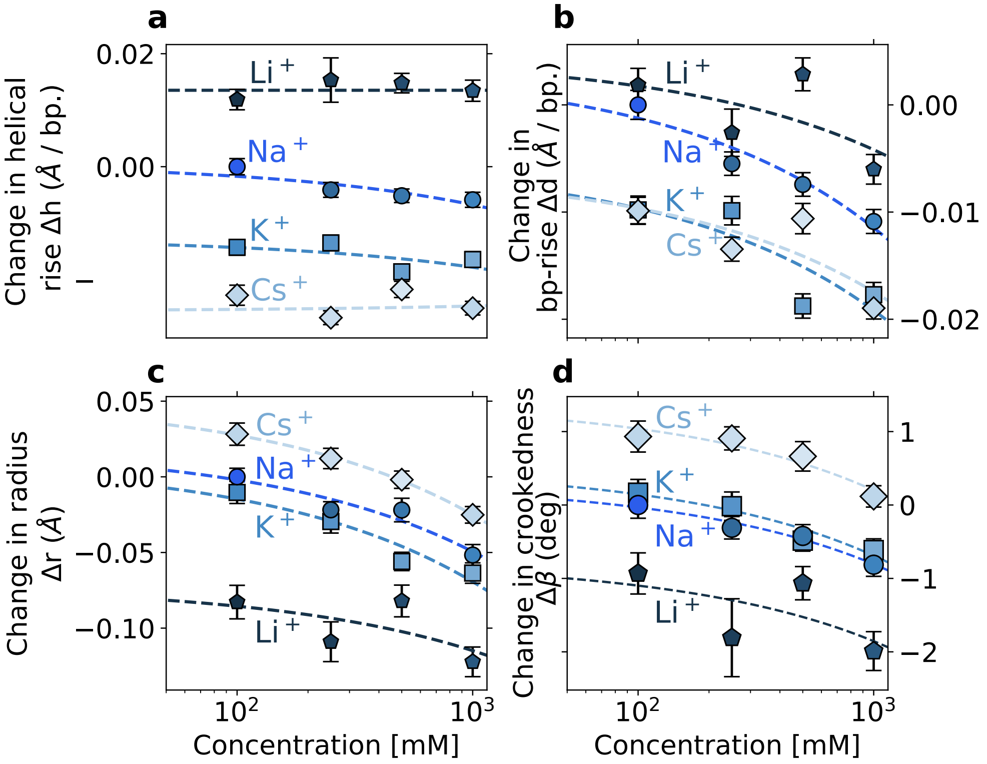


Figure S6: Changes in DNA helical properties as function of monovalent ion type and concentration. (a-d) Changes in the helical rise $\Delta h$, the bp-rise $\Delta d$, the helical radius $\Delta r$, and crookedness $\Delta\beta$, respectively, with respect to 100 mM of KCl. Dashed lines correspond to fittings proportional to the square root of the cation concentration.

#### DNA backbone sub-states in the simulations

As a possible source of change in twist, we evaluate the DNA backbone sub-states, which have been related to changes in the local twist.[S18] In particular, we investigated the sub-states BI and BII defined from the torsion angles $\varepsilon$ and $\zeta$ as BI ($\varepsilon - \zeta < 0$) and BII ($\varepsilon - \zeta > 0$). However, we did not find significant changes in the population of BI/BII sub-states (Figure S7), in agreement with recent results.[S13]

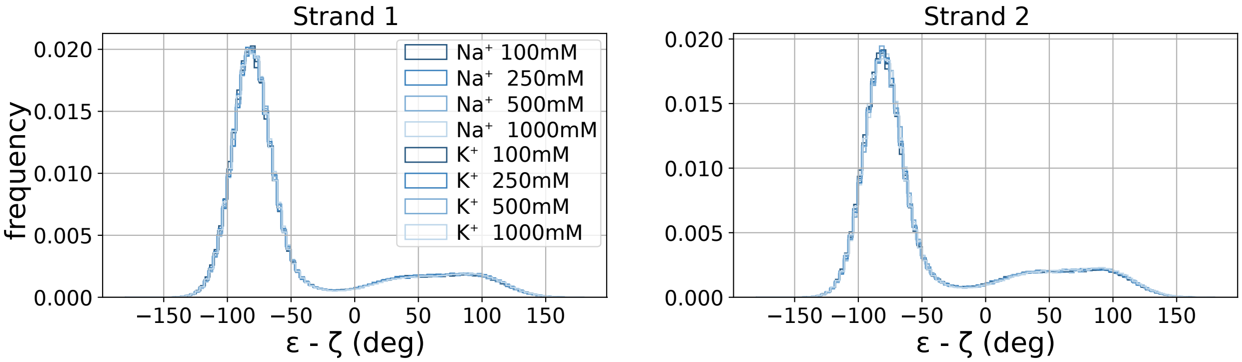


Figure S7: Population of the DNA backbone sub-states BI ($\varepsilon - \zeta < 0$) and BII ($\varepsilon - \zeta > 0$).

#### Changes in twist for divalent cations

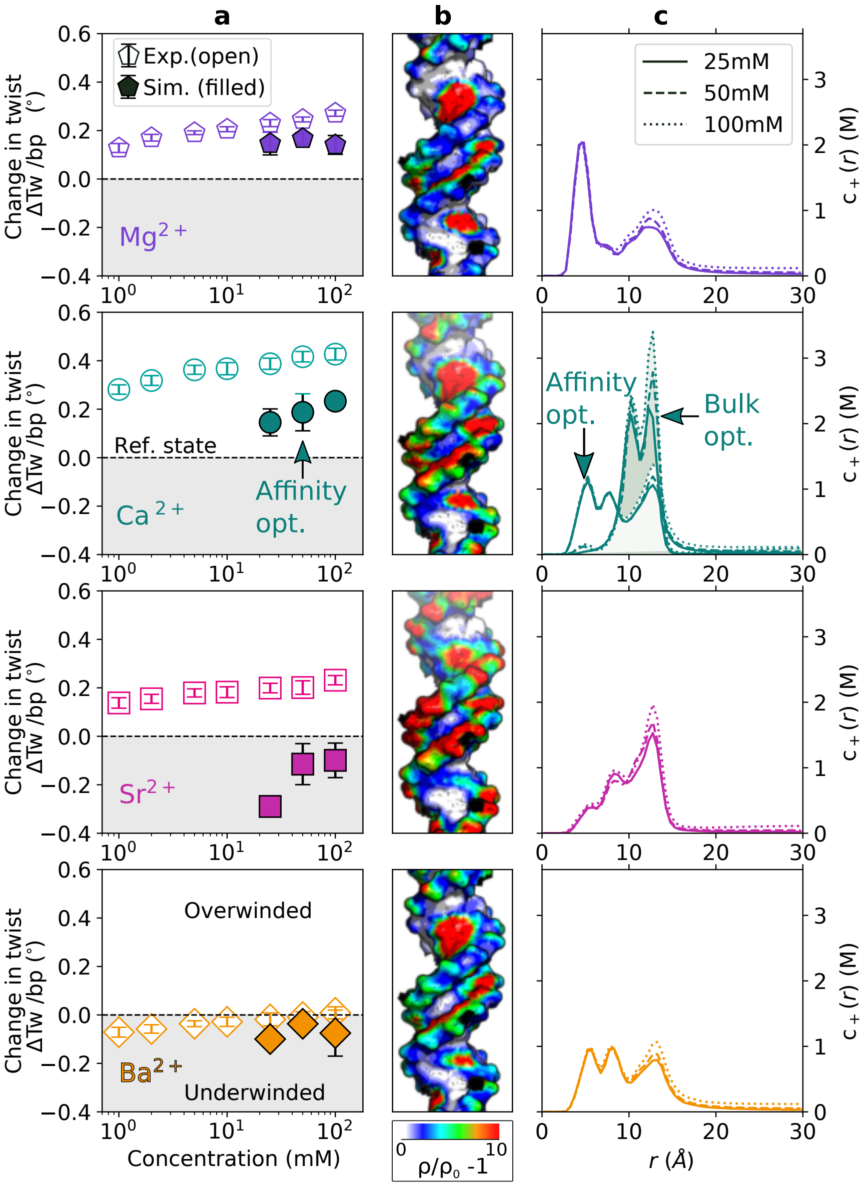


Figure S8: Change in twist for divalent cations. (a) Quantitative comparison of DNA twist obtained from MT experiments (open symbols) and MD simulations (filled symbols) as a function of the ion concentration. (b) Three-dimensional ion distributions. (c) Radial concentration profiles for 25 - 100 mM bulk salt concentration. For $Ca^{2+}$, the concentration profiles for the two force fields[S19,S20] used in this work are shown (see main text for further details).

#### Helical properties for divalent cations

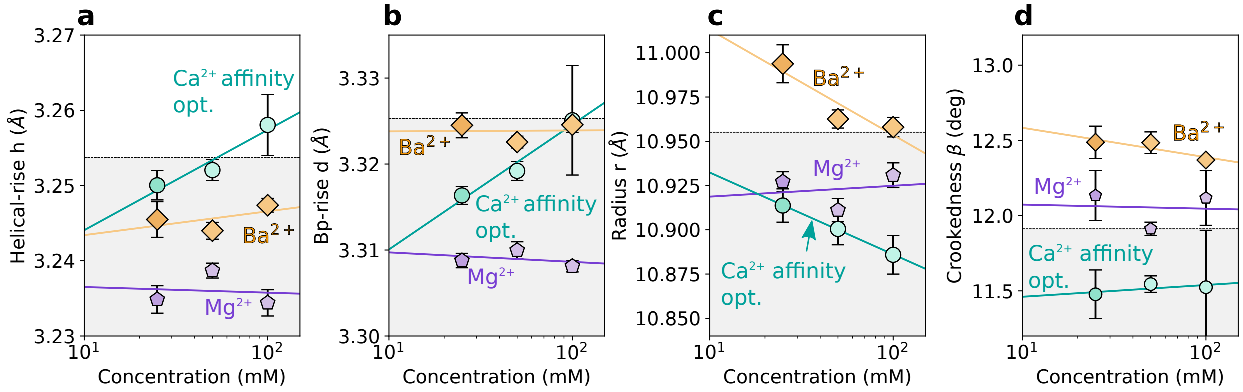


Figure S9: Changes of the characteristic DNA properties for different divalent cations: helical rise h, bp-rise d, radius r, and crookedness $\beta$ for $Mg^{2+}$, $Ca^{2+}$, and $Ba^{2+}$. All results are relative to 100 mM KCl. For $Ca^{2+}$, the data is for the affinity optimized parameter set from Ref.[S20]

### Supplementary Tables

Table S1: Fitting parameters for the experimental change in twist of monovalent cations.

| Ion | $A(c)^B + C$ | | | $A\sqrt{c} + C$ | |
|---|---|---|---|---|---|
| | A | B | C | A | C |
| $Li^+$ | 0.0095 | 0.57 | -0.032 | 0.016 | -0.06 |
| $Na^+$ | 0.012 | 0.49 | -0.2 | 0.011 | -0.2 |
| $K^+$ | 0.017 | 0.44 | -0.13 | 0.011 | -0.11 |
| $Rb^+$ | 0.014 | 0.48 | -0.088 | 0.012 | -0.082 |
| $Cs^+$ | 0.036 | 0.36 | -0.073 | 0.012 | 0.0018 |
| reduced $\chi^2$ | 0.876 | | | 0.97 | |

Table S2: Fitting parameters for the experimental change in twist of divalent cation.

| Ion | $A(c)^B + C$ | | | $A + B \cdot \ln c$ | |
|---|---|---|---|---|---|
| | A | B | C | A | B |
| $Mg^{2+}$ | 0.36 | 0.063 | -0.21 | 0.139 | 0.028 |
| $Ca^{2+}$ | 0.5 | 0.053 | -0.20 | 0.295 | 0.030 |
| $Sr^{2+}$ | 0.42 | 0.04 | -0.27 | 0.142 | 0.018 |
| $Ba^{2+}$ | 0.17 | 0.08 | -0.23 | -0.068 | 0.016 |
| reduced $\chi^2$ | 1.23 | | | 0.84 | |

Table S3: Force field parameters for the metal cations used in this work. The charge and the Lennard-Jones parameters ($\sigma$ and $\epsilon$) are listed. Values were take from Refs.[S19] $\lambda_\sigma^{iR}$ is the optimized scaling factor for the ion–phosphate oxygen interaction as described in Ref.[S20]

| Ion | Charge (e) | $\sigma$ (nm) | $\epsilon$ (kJ mol$^{-}$1) | Charge density (e/nm$^3$) | $\lambda_\sigma^{iR}$ |
|---|---|---|---|---|---|
| $Li^+$ | +1.0 | 0.152 | 0.604 | 67.98 | 1.0 |
| $Na^+$ | +1.0 | 0.261 | 0.115 | 13.43 | 1.0 |
| $K^+$ | +1.0 | 0.299 | 0.604 | 8.93 | 1.0 |
| $Cs^+$ | +1.0 | 0.355 | 0.604 | 5.34 | 1.0 |
| $Mg^{2+}$ | +2.0 | 0.162 | 0.604 | 112.30 | 1.085 |
| $Ca^{2+}$ | +2.0 | 0.226 | 2.339 | 41.36 | 1.027 |
| $Sr^{2+}$ | +2.0 | 0.282 | 1.057 | 21.29 | 1.0 |
| $Ba^{2+}$ | +2.0 | 0.355 | 0.277 | 10.67 | 1.0 |

Table S4: Change in twist ΔTw (deg/bp) for monovalent cations from magnetic tweezer experiments.

| Conc. (mM) | $Li^+$ | | | $Na^+$ | | | $K^+$ | | | $Rb^+$ | | | $Cs^+$ | | |
|---|---|---|---|---|---|---|---|---|---|---|---|---|---|---|---|
| 50 | 0.057 | ± | 0.008 | -0.122 | ± | 0.016 | -0.051 | ± | 0.019 | 0.001 | ± | 0.012 | 0.075 | ± | 0.020 |
| 100 | 0.104 | ± | 0.010 | -0.080 | ± | 0.020 | 0 | | | 0.042 | ± | 0.011 | 0.118 | ± | 0.018 |
| 250 | 0.196 | ± | 0.010 | -0.028 | ± | 0.017 | 0.057 | ± | 0.010 | 0.106 | ± | 0.012 | 0.191 | ± | 0.018 |
| 500 | 0.300 | ± | 0.015 | 0.050 | ± | 0.014 | 0.133 | ± | 0.010 | 0.190 | ± | 0.009 | 0.266 | ± | 0.026 |
| 1000 | 0.473 | ± | 0.023 | 0.149 | ± | 0.012 | 0.227 | ± | 0.015 | 0.299 | ± | 0.014 | 0.361 | ± | 0.018 |

Table S5: Change in twist ΔTw (deg/bp) for divalent cations from magnetic tweezer experiments.

| Conc. (mM) | $Mg^{2+}$ | | | $Ca^{2+}$ | | | $Sr^{2+}$ | | | $Ba^{2+}$ | | |
|---|---|---|---|---|---|---|---|---|---|---|---|---|
| 1 | 0.128 | ± | 0.018 | 0.281 | ± | 0.019 | 0.138 | ± | 0.022 | -0.071 | ± | 0.020 |
| 2 | 0.170 | ± | 0.014 | 0.318 | ± | 0.020 | 0.155 | ± | 0.019 | -0.058 | ± | 0.017 |
| 5 | 0.191 | ± | 0.008 | 0.362 | ± | 0.019 | 0.179 | ± | 0.017 | -0.036 | ± | 0.012 |
| 10 | 0.206 | ± | 0.010 | 0.367 | ± | 0.025 | 0.182 | ± | 0.022 | -0.028 | ± | 0.019 |
| 25 | 0.231 | ± | 0.014 | 0.387 | ± | 0.023 | 0.199 | ± | 0.020 | -0.019 | ± | 0.027 |
| 50 | 0.246 | ± | 0.010 | 0.416 | ± | 0.022 | 0.201 | ± | 0.028 | -0.009 | ± | 0.022 |
| 100 | 0.272 | ± | 0.013 | 0.427 | ± | 0.024 | 0.232 | ± | 0.019 | 0.009 | ± | 0.025 |
